# Dysregulated intestinal nutrient absorption in obesity is associated with epigenomic alterations in epithelia

**DOI:** 10.1101/2024.05.06.591758

**Authors:** Dilhana S. Badurdeen, Zhen Li, Jeong-Heon Lee, Tao Ma, Aditya Vijay Bhagwate, Rachel Latanich, Arjit Dogiparthi, Tamas Ordog, Olga Kovbasnjuk, Vivek Kumbhari, Jennifer Foulke-Abel

## Abstract

Obesity is an epidemic with myriad health effects, but little is understood regarding individual obese phenotypes and how they may respond to therapy. Epigenetic changes associated with obesity have been detected in blood, liver, pancreas, and adipose tissues. Previous work using human organoids found that dietary glucose hyperabsorption is a steadfast trait in cultures derived from some obese subjects, but detailed transcriptional or epigenomic features of the intestinal epithelia associated with this persistent phenotype are unknown. This study evaluated differentially expressed genes and relative chromatin accessibility in intestinal organoids established from donors classified as non-obese, obese, or obese hyperabsorptive by body mass index and glucose transport assays. Transcriptomic analysis indicated that obese hyperabsorptive subject organoids have significantly upregulated dietary nutrient absorption transcripts and downregulated type I interferon targets. Chromatin accessibility and transcription factor footprinting predicted that enhanced HNF4G binding may promote the obese hyperabsorption phenotype. Quantitative RT-PCR assessment in organoids representing a larger subject cohort suggested that intestinal epithelial expression of CUBN, GIP, SLC5A11, and SLC2A5 were highly correlated with hyperabsorption. Thus, the obese hyperabsorption phenotype was characterized by transcriptional changes that support increased nutrient uptake by intestinal epithelia, potentially driven by differentially accessible chromatin. Recognizing unique intestinal phenotypes in obesity provides a new perspective in considering therapeutic targets and options to manage the disease.

## INTRODUCTION

Obesity is a growing global public health crisis, with over 1.9 billion adults being overweight, and 650 million obese.(1) In 2016, the annual healthcare cost of adult obesity in the U.S. was $260.6 billion.(2) The World Health Organization recognizes obesity as a risk factor for type 2 diabetes, coronary heart disease, asthma, stroke, and multiple cancers. Obesity-associated diseases place an additional burden on the healthcare system, with medical costs estimated to be 30% higher for obese individuals (BMI ≥30) compared to those that are lean (BMI ≤25). Among the available therapies, bariatric surgery is the most effective intervention for significant and sustained weight loss.(3) However, surgical outcomes are heterogeneous with 15-35% of patients experiencing weight loss failure (<50% excess weight loss at 12-18 months) after roux-en-Y gastric bypass (RYGB).(4, 5) Numerous hypotheses have fallen short of fully identifying and characterizing molecular changes catalyzed by bariatric procedures.(6) Prediction of individualized weight loss success based on phenotypic classification would be important to not only guide selection of an effective intervention, but also to minimize potential risks and complications from ineffective procedures.

Among the many phenotypes associated with obesity, accelerated intestinal glucose absorption observed in nondiabetic obese subjects(7) is a striking example of an altered physiological response that likely exacerbates the disease yet lacks full mechanistic understanding. Intestinal organoids/enteroids are a unique experimental tool that stably retain intestinal region- and donor-specific genotype and phenotypes.(8) In our prior study using patient-derived jejunal organoid monolayers from non-obese (control group, G1) and obese subjects,(9) we found that a subset of obese donors exhibited significantly increased epithelial glucose absorption capacity (obese hyperabsorption group, G2) relative to G1 non-obese controls or other obese donor organoids (G3). This observation suggested that intestinal metabolic phenotypes may be genetically encoded and differ among obese subjects. It further intimated that genetic regulation of dietary sugar hyperabsorption may contribute to metabolic disease in G2 subjects. Considering the relatively recent global surge in obesity development and the paucity of evidence for specific protein-coding gene mutations to drive this outcome, we hypothesized that potential stable epigenetic modifications in G2 individuals may account for the hyperabsorptive phenotype. In this study, we analyzed the transcriptomes of differentiated organoid monolayers derived from 3 representative donors for each group (G1, G2, and G3) to identify differentially expressed genes characteristic of each group. We also conducted an assay for transposase-accessible chromatin sequencing (ATAC-seq) to determine differential chromatin accessibility, and then carried out ATAC footprinting(10) and motif analysis to detect putative transcription factors differentially occupying chromatin among the three groups. Transcript levels distinctive for differing metabolic profiles were additionally validated in differentiated organoid monolayers derived from an expanded cohort of non-obese and obese subjects. Collectively, our data provides important insights into the dysregulated transcriptional state and altered open chromatin landscape of intestinal epithelia in obese glucose hyperabsorptive subjects.

## RESULTS

### Differential enrichment of metabolism-associated transcripts in obese donor-derived differentiated organoid monolayers exhibiting glucose hyperabsorption

Donor organoid lines were previously characterized as non-obese controls (G1, BMI ≤29.9, n=3), obese with elevated intestinal glucose absorption relative to non-obese controls (G2, BMI ≥35, n=3), or obese with intestinal glucose absorption similar to non-obese controls (G3, BMI ≥35, n=3) (**Table 1**).(9) Bulk RNA-seq of differentiated organoid monolayers representing 3 donors in each group was conducted to identify differentially expressed genes (DEG, absolute log_2_ fold change>2, *p*<0.05) (**Fig 1A** and **Supplementary Table S1**). Principal component analysis demonstrated that the overall transcriptional profiles among the three groups have considerable overlap, and that one individual in each group appeared to segregate slightly from their respective group (**Fig 1B**). Comparison of G2 to either G1 or G3 each revealed 456 DEGs, many of which function in dietary nutrient absorption, metabolism, and regulation (**Fig 1C, D**). In contrast, G1 and G3 were distinguished by only 27 DEGs, suggesting highly similar transcriptomes (**Fig 1E**). The enriched gene ontology biological processes (GOBP) in G2 included glucose transport, cholesterol transport, and triglyceride homeostasis (Cluster #1, **Fig 1F, G**). Downregulated GOBP in G2 epithelia included type I interferon signaling associated with regulation of innate immune responses (Cluster #2, **Fig 1F, G**).

**Figure 1.**
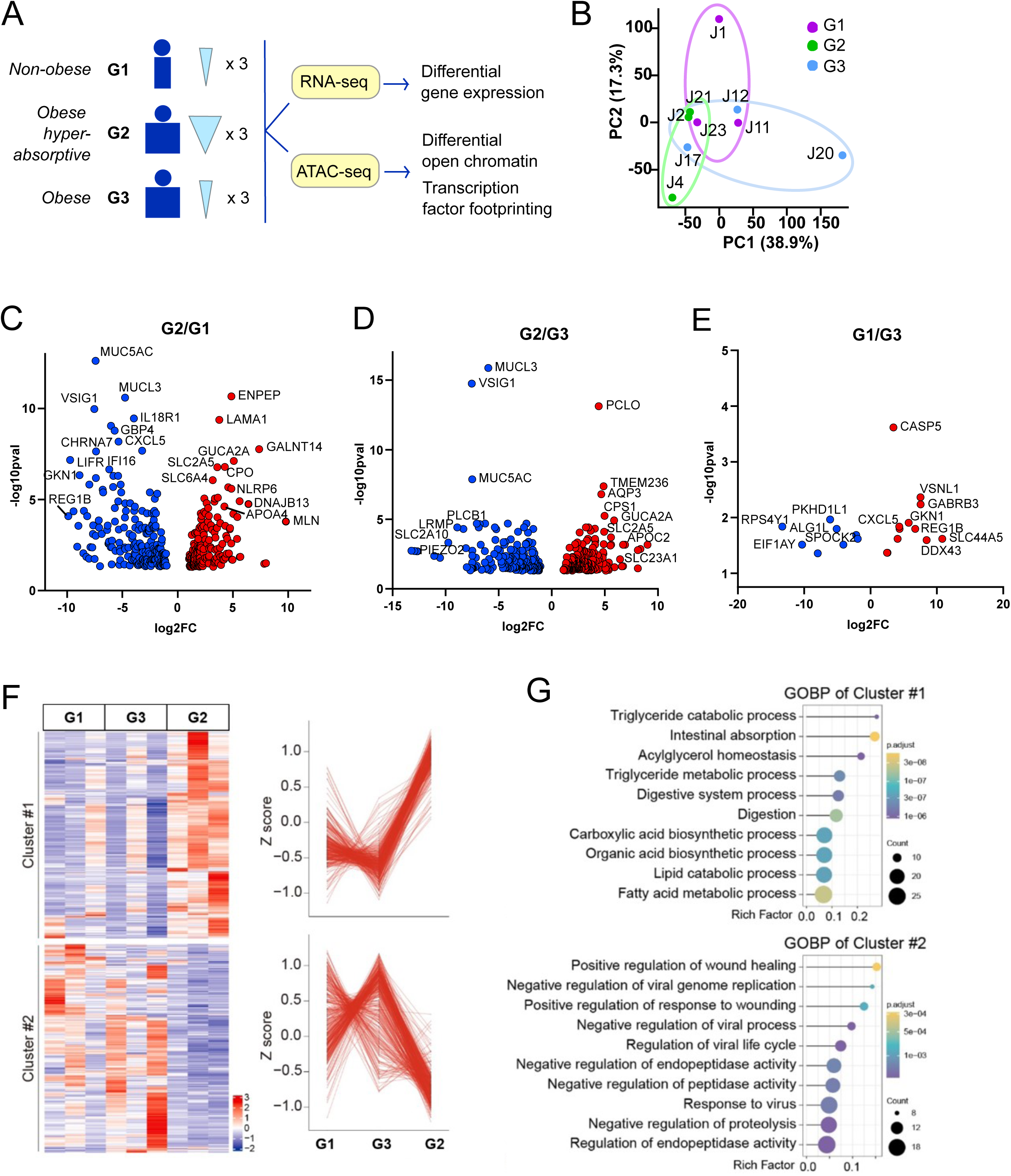
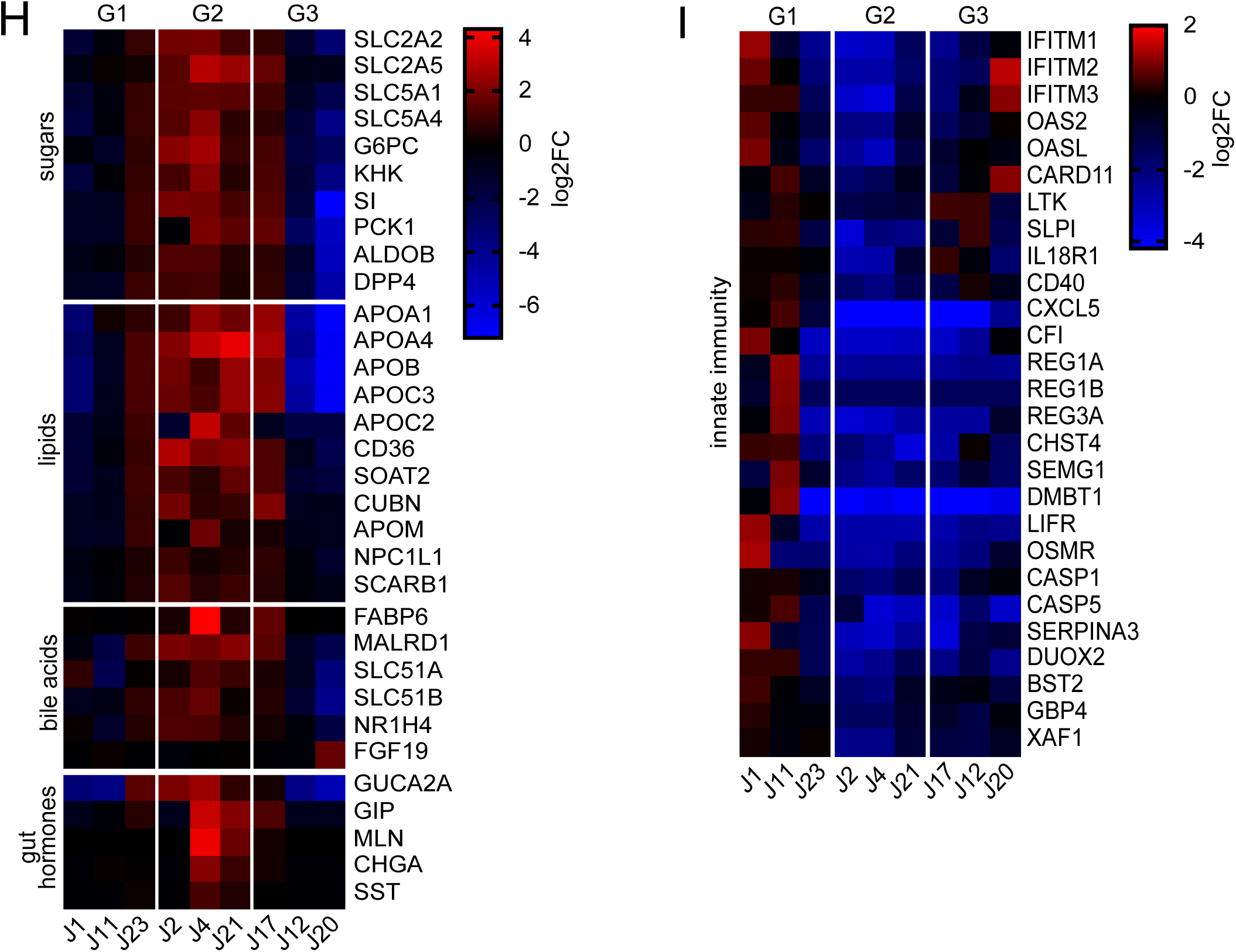
Differential gene expression among the organoid donor groups. (**A**) Experimental design and donor phenotypes of groups 1, 2, and 3. (**B**) Principal component analysis (PCA) of the nine donors following RNA-seq analysis. Colors distinguish donors in each of the three groups. (**C**) Volcano plot depicting significantly upregulated (red) or downregulated (blue) transcripts (log_2_FC > |2|, *p* < 0.05) in paired comparisons of G2/G1, (**D**) G2/G3, and (**E**) G1/G3. (**F**) Heatmap of relative expression for significant DEGs in G2 compared to G1 or G3. Cluster 1 contains DEG that were significantly upregulated in G2, and Cluster 2 contains DEG that were significantly downregulated in G2. Z scores for each gene were computed from log_2_(FPKM+1) contained in the respective clusters. (**G**) Gene ontology (GO) terms representing major biological processes characteristic of Clusters 1 and 2. (**H**) Heatmap (log_2_FC) for selected transcripts distinctively upregulated in G2 that are associated with dietary sugar transport/metabolism, fat absorption/transport, bile acid absorption, and gut hormones. (**I**) Heatmap of selected transcripts distinctively downregulated in G2 that loosely group as factors involved in innate immune functions.

**Table 1.** Organoid donor characteristics. Deidentified subject designator (organoid line), group classification (G1, non-obese; G2, obese hyperabsorptive; G3, obese) based on glucose transport phenotype, demographic information (age, sex), body mass index (BMI), and passage number for each organoid line studied. n.r., not recorded; n.a., not applicable.

| Group | Organoid Line | Originating Institution | Absorption Phenotype | Age | Sex | BMI | Organoid passage used in RNA-based experiments | Patient ID previously reported(9) |
| --- | --- | --- | --- | --- | --- | --- | --- | --- |
| G1 | J1 | Johns Hopkins | low | n.r. | F | ≤25 | P38 | 3 |
| G1 | J11 | Baylor | low | n.r. | F | 23 | P29 | 4 |
| G1 | J23 | Johns Hopkins | low | 46 | F | 21 | P11 | 5 |
| G2 | J2 | Baylor | high | 52 | F | ≥35 | P81 | 7 |
| G2 | J4 | Baylor | high | n.r. | F | ≥35 | P33 | 8 |
| G2 | J21 | Johns Hopkins | high | 42 | F | 43 | P10 | 9 |
| G3 | J12 | Baylor | low | n.r. | M | ≥35 | P28 | 13 |
| G3 | J17 | Johns Hopkins | low | 49 | F | 53 | P15 | 16 |
| G3 | J20 | Johns Hopkins | low | 48 | F | 51 | P13 | 14 |
| G1 | 1 | Johns Hopkins | low | 81 | F | 22.8 | P16 | n.a. |
| G1 | 23 | Johns Hopkins | low | 57 | M | 23 | P9 | n.a. |
| G1 | 24 | Johns Hopkins | low | 63 | F | 28 | P7 | n.a. |
| G1 | 25 | Johns Hopkins | low | 63 | M | 28.2 | P8 | n.a. |
| G1 | 26 | Johns Hopkins | low | 65 | F | 24 | P6 | n.a. |
| G1 | 27 | Johns Hopkins | low | 73 | F | 23.8 | P4 | n.a. |
| G1 | 28 | Johns Hopkins | low | 67 | F | 27.1 | P5 | n.a. |
| G1 | 0020 | Mayo | low | 20 | M | 23.1 | P12 | n.a. |
| G1 | 0021 | Mayo | low | 21 | F | 22.1 | P10 | n.a. |
| G1 | 0022 | Mayo | low | 37 | M | 19.5 | P10 | n.a. |
| G2 | 0003 | Mayo | high | 51 | F | 32.7 | P7 | n.a. |
| G2 | 0004 | Mayo | high | 58 | M | 36.3 | P5 | n.a. |
| G2 | 0005 | Mayo | high | 73 | F | 36.3 | P8 | n.a. |
| G2 | 0009 | Mayo | high | 73 | F | 36.7 | P6 | n.a. |
| G2 | 0012 | Mayo | high | 55 | F | 34.3 | P4 | n.a. |
| G2 | 0013 | Mayo | high | 32 | F | 42.7 | P6 | n.a. |
| G2 | 0014 | Mayo | high | 47 | F | 36.9 | P7 | n.a. |
| G2 | 0015 | Mayo | high | 46 | F | 43.5 | P4 | n.a. |
| G2 | 0018 | Mayo | high | 71 | F | 38.3 | P3 | n.a. |
| G2 | 0031 | Mayo | high | 55 | F | 32.1 | P4 | n.a. |

#### Sugar transport and metabolism

Significantly increased transcripts in G2 included the major facilitators of intestinal glucose and fructose absorption: SLC2A2, SLC2A5, SLC5A1, and SLC5A4 (**Fig 1H**), which confirmed and expanded the findings of prior work.(9) As seen previously, levels of gluconeogenic enzymes glucose-6-phosphatase (G6PC) and phosphoenolpyruvate carboxykinase (PCK1) were also elevated. Ketohexokinase (KHK) and aldolase B (ALDOB) were also increased in G2, suggesting that the epithelia is primed to catabolize an influx of dietary fructose entering through GLUT5. Elevated transcripts for the brush border enzyme sucrase-isomaltase (SI) may enhance digestion of sucrose and maltose into simple sugars for intestinal uptake, and dipeptidyl peptidase IV (DPP4) would have potential to activate gut hormones involved in metabolic regulation.

#### Lipid transport and metabolism

Dietary fatty acids can be either passively absorbed or transit via membrane-bound proteins. Transporters situated at the apical plasma membrane including fatty acid translocase (CD36) and apolipoprotein M (APOM) were significantly upregulated in G2 (**Fig 1H**), suggesting enhanced uptake ability. Additional fatty acid carriers included apolipoproteins APOA1, APOA4, APOB, APOC2, and APOC3 that participate in lipid transport and act as enzyme cofactors for lipid metabolism. Other amplified transcripts of note included receptors for lipoproteins (CUBN and SCARB1), a channel for cholesterol-containing micelle import (NPC1L1), and SOAT2, which catalyzes cholesterol esterification necessary for lipoprotein assembly. The rate limiting enzyme in mitochondrial fatty acid oxidation (CPT1A) was relatively unchanged, suggesting that enterocytes are primarily facilitating fatty acid transport rather than executing catabolism.

#### Bile acid absorption and signaling

Bile acids are normally introduced in the duodenum to assist with solubilizing dietary fats and are subsequently reabsorbed in the distal ileum. Surprisingly, G2 demonstrated transcriptional enrichment in the jejunum for proteins involved in bile acid reabsorption that are usually most prominent in distal ileum (**Fig 1H**). These included the major intracellular carrier for bile acids, fatty acid binding protein 6 (FABP6), and the basolateral bile acid exporter subunits SLC51A and SLC51B, as well as the bile acid responsive transcription factor FXR (NR1H4) and MALRD1, a cofactor known to augment intestinal FGF19 secretion.(^11^) FGF19 transcripts were similar among the three groups, which would be expected due to the absence of exogenous bile acids in the organoid culture media to induce expression.

#### Intestinal hormones

Peptide hormones were another class of signaling molecule with differential expression that distinguished G2 from G1 and G3. G2 subjects were found to overexpress guanylin (GUCA2A), an intestinal peptide hormone produced by enterocytes and Paneth cells with pleiotropic effects on epithelial transport via induction of cGMP synthesis (**Fig 1H**). Upregulated transcripts related to enteroendocrine cell (EEC)-specific hormones, including glucose-dependent insulinotropic polypeptide (GIP), motilin (MLN), somatostatin (SST), and the prohormone chromogranin A (CHGA) were also detected. Given the relative rarity of EECs in organoids grown under conditions that do not optimize EEC development,(12) the intrinsic emergence of these hormone transcripts is further evidence of differential transcriptional controls in the G2 state.

Collectively, these results indicated that the obese glucose hyperabsorption phenotype (G2) is characterized by upregulated transcription of nutrient transporters and metabolic regulators that may contribute to dysregulated energy homeostasis in a subset of obese subjects.

### Downregulation of epithelial innate immune response transcripts in G2 and G3

Many of the downregulated targets distinctive in G2 included interferon-stimulated genes (ISGs) and downstream molecules linked to the type I interferon (IFN) signaling pathway (**Fig 1I**). These included antiviral components such as interferon induced transmembrane proteins IFITM1, IFITM2, and IFITM3, which restrict viral entry, and OAS2 and OASL, which act as sensors for double-stranded RNA. Innate defense molecules including antimicrobial proteins REG1A, REG1B, REG3A, SLPI, and SEMG1 were also significantly decreased relative to G1. Receptors with immune-associated functions including IL18R1, LIFR, OSMR, CD40, and LTK were also reduced. Interestingly, G3 shared downregulation of many innate immune-related genes that were observed in G2, implying that innate immune repression may be shared among obese subjects regardless of their dietary nutrient absorption phenotype.

### Differential open chromatin regions in G2 suggest potential epigenetic alterations

ATAC-seq was performed using the same differentiated organoid monolayer groups to investigate the potential for an altered chromatin landscape that would account for the marked differential gene expression in G2 relative to G1 or G3 (**Supplementary Table S2**). Comparison of G2 to G1 yielded 2,051 differential sites (**Fig 2A**), and G2 vs G3 indicated 3,549 differential sites (**Fig 2B**). G1 and G3 differed by a more modest 491 sites (**Fig 2C**). The categorical locations of differential open chromatin were similarly distributed in each of the three paired group comparisons. Notably, there were distinct open chromatin regions proximal to transcriptional start sites (TSS) in G2 that were not equivalent in G1 nor G3 (**Fig 2D**). Many of these sites coincided with increased levels of downstream gene transcripts, suggesting an increase in enhancer activity.

**Figure 2.**
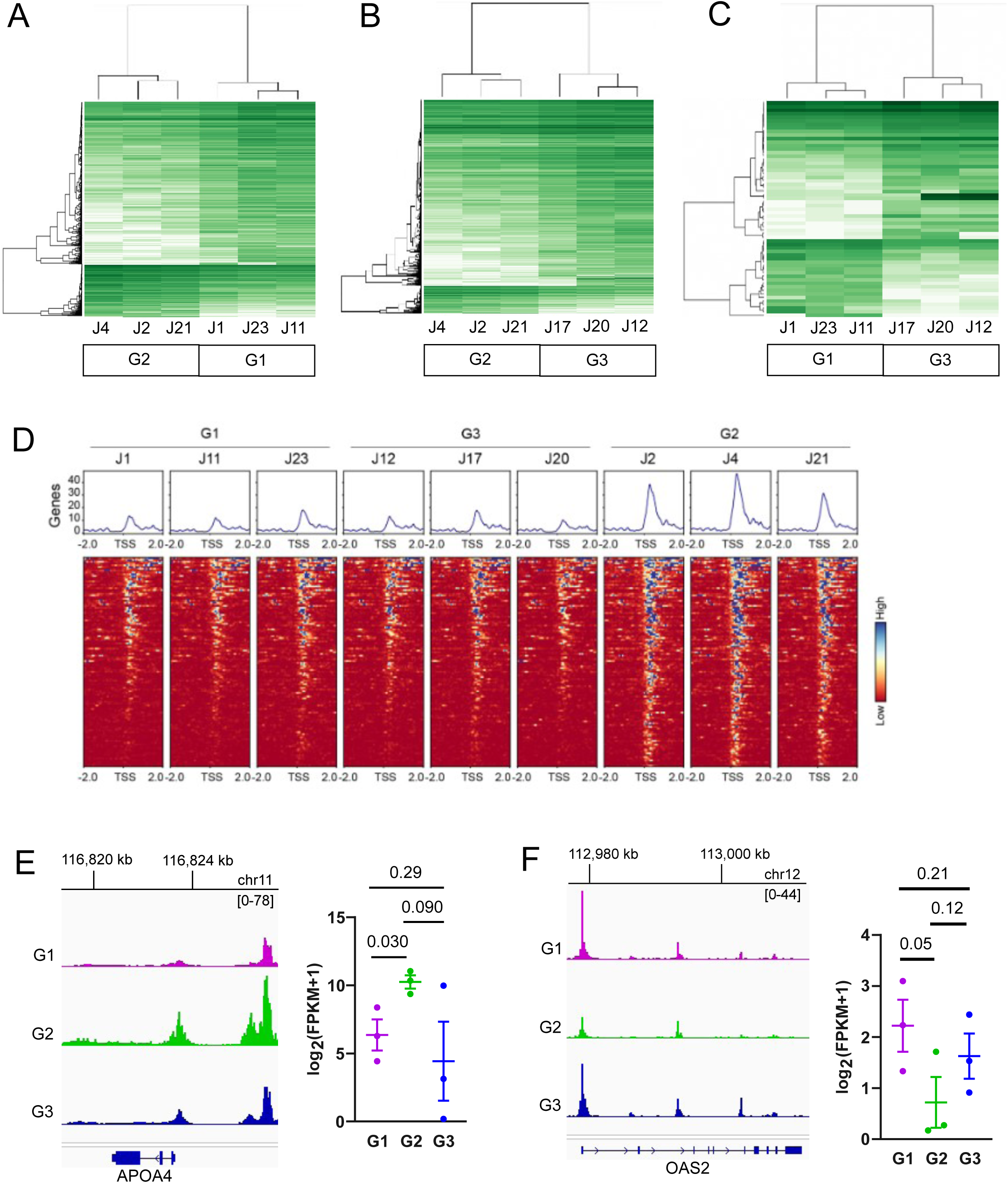
Differential open chromatin regions among the organoid donor groups. (**A**) Differential open chromatin regions (log_2_FC > |2|, *p* < 0.05) identified from ATAC-seq in G1 vs G2, (**B**) G2 vs G3, and (**C**) G1 vs G3. (**D**) Heatmaps of differential open chromatin proximal to transcription start sites (TSS, within 2 kb upstream or downstream) for each donor. (**E**) Summed ATAC-seq peak tracks for each of the three groups representing the upstream and protein coding region for APOA4, which had increased accessibility in G2. (**F**) Summed peak tracks for OAS2, which had decreased accessibility in G2 relative to the other groups.

When integrated with transcriptional data, pathways uniquely enriched in G2 were centered on digestion and lipid transport/metabolism, an example of which is APOA4 (**Fig 2E**). In contrast, innate immune-related transcripts appeared to be less prominent due to less open chromatin near the promoter region of genes such as interferon-inducible antiviral enzyme 2’-5’-oligoadenylate synthetase 2 (OAS2) (**Fig 2F**).

### ATAC footprinting indicates differential binding of transcription factors (TFs) associated with metabolic and innate immune regulation in G2

ATAC footprinting in G2 compared to G1 (**Fig 3A**) or G3 (**Fig 3B**) yielded significant differential binding predictions that were in general agreement with transcriptional analyses and were highly similar between the two comparators. Comparison of G1 and G3 did not render any significant differential binding results, which further reflects their transcriptional similarity. G2 footprints suggested increased binding of nuclear receptor ligand-activated TFs known to regulate digestive and metabolic processes, including hepatocyte nuclear factor 4 paralogs (HNF4A, HNF4G) and numerous retinoic acid X receptor (RXR) heterodimers. Lower scoring but significant binding prediction also implicated peroxisome proliferator activated receptors (PPARA, PPARD, PPARG) which function as key regulators of lipid metabolism.

**Figure 3.**
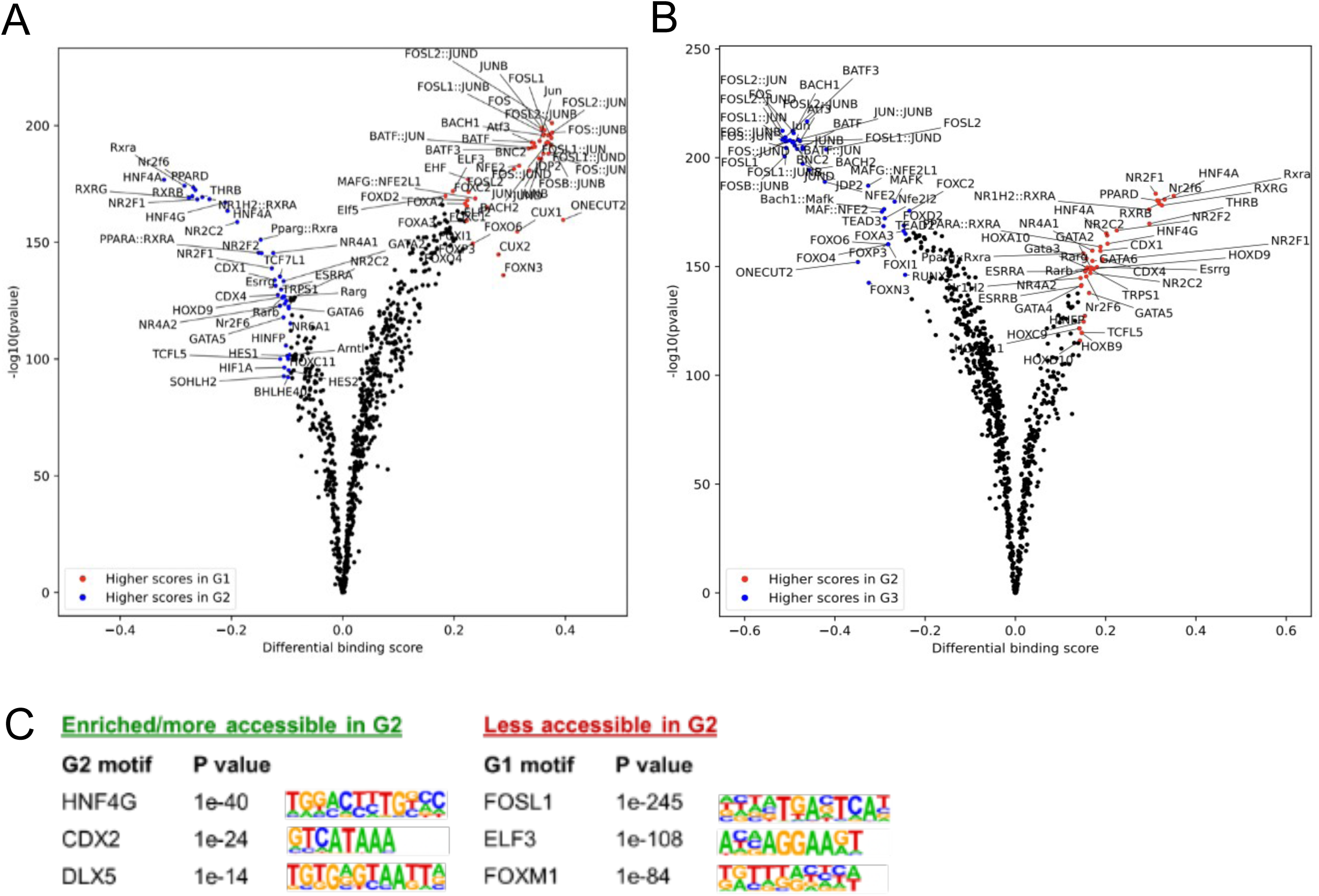
Differential transcription factor binding events inferred from ATAC footprinting. (**A**) Volcano plot of differential binding scores in G1 vs G2 and (**B**) G2 vs G3. There were no significant differential binding events in G1 vs G3. (**C**) Top three most and least accessible TF motifs in G2 when compared to G1, ranked by *p*-value.

Integrated binding motif analysis ranked HNF4G as the top factor involved in G2 upregulated gene transcription (*p* = 1e-40) (**Fig 3C**). HNF4G transcripts were also slightly upregulated (log_2_FC = 1.15) in G2, suggesting that it may exert dominance in transcriptional regulation in the G2 state. Further, RNA-seq FPKM values for HNF4A and HNF4G were approximately equivalent in G1 and G3, but HNF4G FPKM was twice as high in G2 as in G1 or G3. A combination of increased open chromatin sites conducive to HNF4 paralog binding and increased HNF4G levels suggest that the G2 state may be driven at least in part by HNF4G activity.

Decreased footprinting was observed for Jun proto-oncogene (JUN), Fos proto-oncogene (FOS), and activating transcription factor (ATF) family proteins that constitute the activator protein-1 (AP-1) TF complex. AP-1 complexes regulate diverse cell responses related to inflammation, proliferation, differentiation, and apoptosis. Motif analysis ranked FOSL1 sites as least accessible in G2 compared to G1 (*p* = 1e-245) or to G3 (*p* = 1e-427). Although there were no significant differential footprints detected in G1 versus G3, motif analysis did indicate that FOSL2::JUN sites were less accessible in G3 (*p* = 1e-20), hinting that some AP-1 mediated processes may be attenuated in obese subjects with normative dietary nutrient absorption profiles. This was corroborated in the RNA-seq analysis showing that some of the innate immune-associated transcripts notably downregulated in G2 were also decreased in G3 relative to G1 (**Fig 1I**).

### Analysis of differentiated organoid monolayers from additional subjects corroborates upregulated genes characteristic of G2

To follow up on the general findings from the nine selected donor organoids analyzed in RNA-seq and ATAC-seq experiments, we conducted glucose transport assays in differentiated organoid monolayers to identify additional G1 and G2 subjects (n=10 subjects/group, **Fig 4A**). This was followed by quantitative real-time PCR (qRT-PCR) analysis of cDNA isolated from differentiated monolayers. A panel of 10 genes related to dietary sugar/fat metabolism and gut hormones was chosen to validate transcriptional trends observed by RNA-seq. This expanded group of organoids did not identify any additional G3 donors. As expected, organoids that were phenotyped as G2 in glucose transport assays exhibited markedly increased transcript levels for the panel genes (presented quantitatively as ΔC_T_ values) relative to G1 donor organoids that were generally consistent with the RNA-seq findings (**Fig 4B, C**). Individual genes were compared among larger groups of G1 and G2 donor organoids using receiver-operator curves (ROC), and the calculated area under the curve (AUC) indicated that some transcripts (GIP, CUBN, SLC5A11, SLC2A5) appeared to have high predictive value (AUC≥0.910, *p*≤0.002), while other transcripts (GUCA2A, APOA4) were less robust indicators of group association (AUC≤0.780, p≤0.03) (**Fig 4D**).

**Figure 4.**
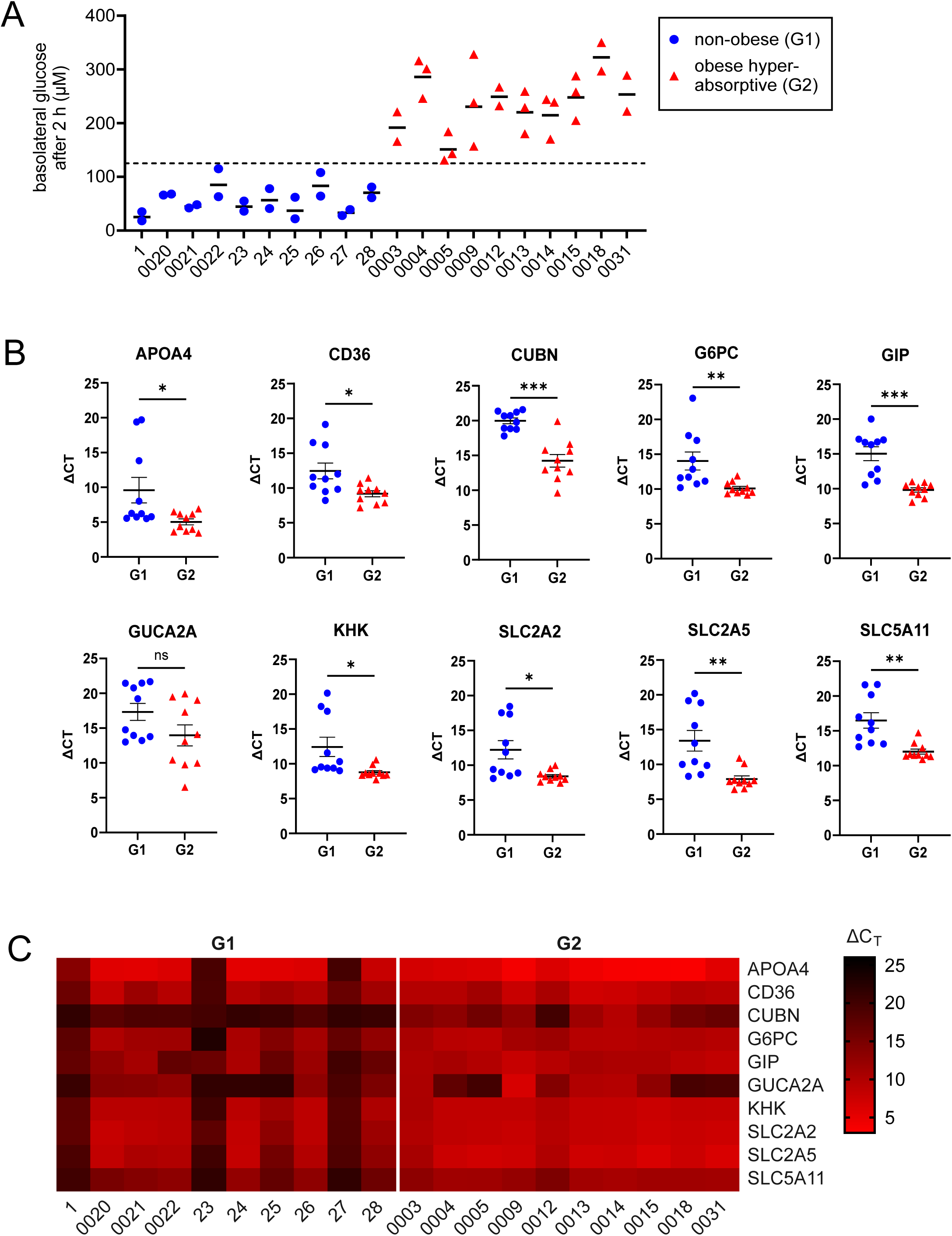
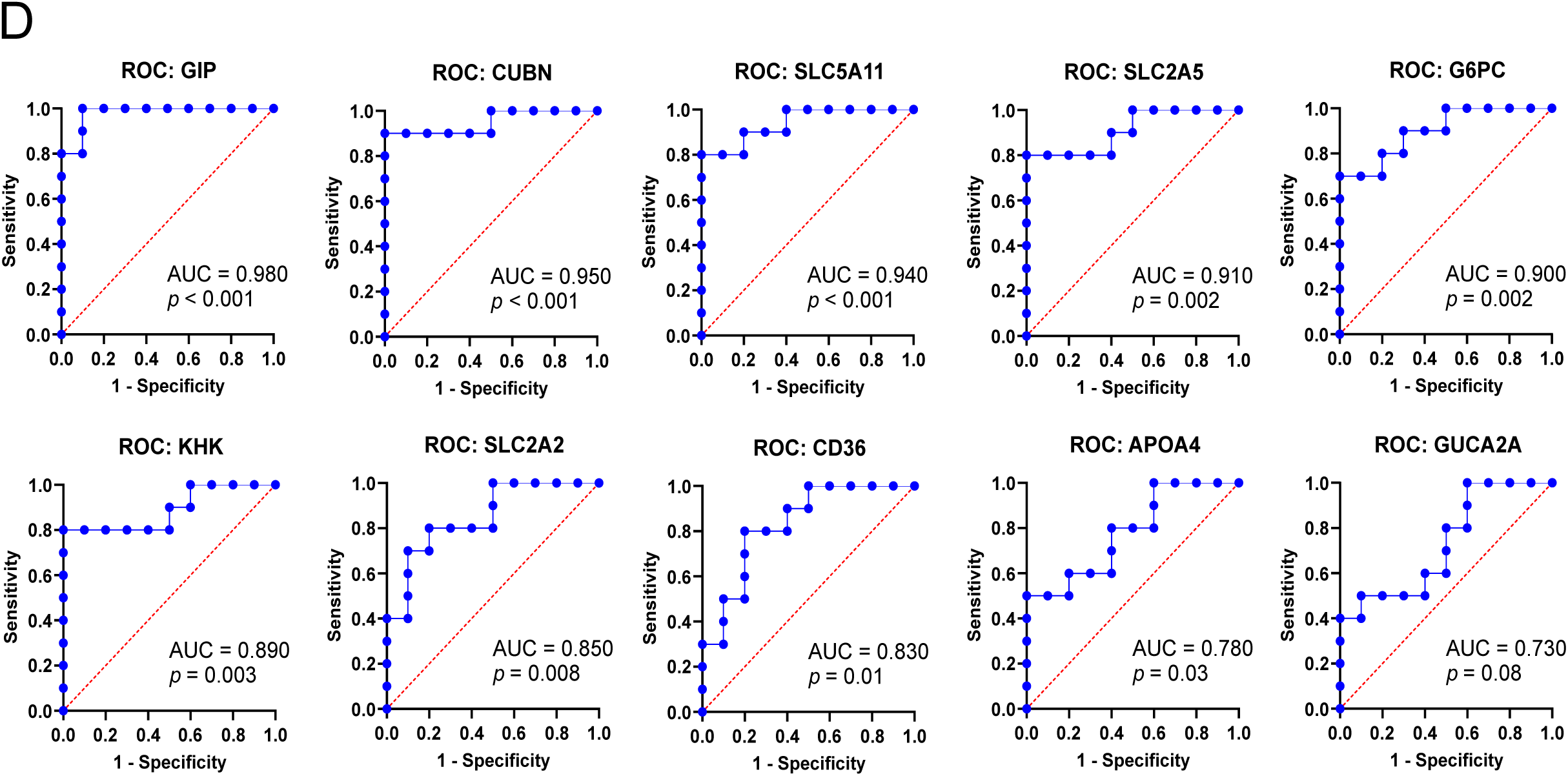
Validation of differentially expressed intestinal epithelial biomarkers in obese hyperabsorptive subjects. **(A)** Glucose transport measured as the D-glucose concentration accumulated in the lower Transwell chamber (basolateral buffer) after 2 h feeding with buffer containing 25 mM D-glucose. Organoids were established from non-obese or obese subjects (n=10 organoids in each group). Each dot represents an independent transport experiment and is the average of two Transwell technical replicates. *, p≤0.05; **, p≤0.01; ***, p≤0.001. Absorption phenotype/group assignment was determined using a cutoff of 125 μM D-glucose (dotted line). Organoids were unique from those analyzed by next-generation sequencing and the prior Hasan et al study. (**B**) Transcripts measured by quantitative real-time PCR (qRT-PCR) expressed as ΔC_T_, the difference between the target gene C_T_ and the average of the C_T_ for housekeeping transcripts RN18S and GAPDH. (**C**) Heatmap of ΔC_T_ values depicted in panel B to illustrate individual donor organoid ΔC_T_ trends. Increasing intensity of the red color indicates a smaller ΔC_T_ value (higher transcript levels). (**D**) Receiver-operator curves (ROC) for each biomarker analyzed by qRT-PCR. Each graph includes the respective area under the curve (AUC) and *p*-value.

## DISCUSSION

Intestinal organoids derived from individuals with metabolic syndrome indicate that some subjects have distinct transcriptional profiles consistent with dysregulated energy balance. The intestinal stem cells of G2 subjects appeared to maintain genetic memory of their in vivo state with entrained upregulation of fatty acid uptake, dietary sugar transport and metabolism, and specific gut hormones that was independent of their immediate diet, given that all of the organoids were provided with the same media/nutrition throughout the experimental process. Additionally, the transcriptional differences observed were intrinsic to the epithelia and not dependent on other tissue components (mesenchyme, immune, muscle, endothelia, lymphatics, neurons, or microbes). The transcriptional signatures appear stable in organoid culture, as the cell lines tested were from a range of passages between 10-81 and have been cryopreserved and rethawed multiple times.

Diet may play a major role in catalyzing the intestinal adaptation toward a hyperabsorptive phenotype. Gut epithelia have the ability to undergo a rapid transcriptional shift toward increased fatty acid absorption capacity in response to dietary nutrient influx, even observed within days of transitioning mice from standard chow to a high fat diet.(13) Numerous studies have examined the influence of diet composition on intestinal stem cell (ISC) behavior in animal models, finding that obesogenic high sugar and/or high fat diets alter epithelial proliferation and lineage differentiation trajectories, and confirming that ISCs can sense and respond to nutrients.(14–16) Studies evaluating the effect of diet on ISCs could also explain the altered gut hormones evident in G2 subjects. Increased levels of GIP, SST, and MLN in G2 subjects are consistent with observations that dietary exposures alter development of enteroendocrine cell subtypes in mice.(13, 14) However, the duration and magnitude of dietary imbalance that may lead to durable epigenetic modifications is not well defined. This study provides insight not only toward the contribution of intestinal epithelia as a facilitator of excess nutrient absorption but also suggests that acquisition of hard-wired transcriptional changes in these functions may undermine efforts to restrict calorie intake.

TF footprinting and DNA binding motif analysis implicated HNF4 as the top-scoring regulator of the G2 hyperabsorptive phenotype, and the role of HNF4 in intestinal absorptive/metabolic function is strongly supported by prior animal studies. HNF4 has become known for orchestrating critical programming of intestinal enterocyte maturation, brush border function, and fatty acid oxidation.(17–20) HNF4A and HNF4G function as highly redundant paralogs in the intestinal epithelium, with HNF4G being predominantly intestine-restricted.(21) Hnf4g^-/-^ mice were protected from diet-induced obesity, likely due to intestinal lipid malabsorption and increased GLP-1 levels.(22) Thus, the HNF4 paralogs and origin of altered intestinal binding site accessibility in human obesity will be important targets for future study. Increased occupancy of PPARD and retinoic acid X receptor α (RXRA) binding sites were also predicted in G2, consistent with known roles in sensing and responding to diet. Despite evidence of more active HNF4, PPARD, and RXRA, the transcripts for rate-limiting enzymes involved in fatty acid oxidation (FAO) such as CPT1A were not upregulated. This could be a consequence of the limited fatty acid content in standard organoid medium, so it would be interesting to assess whether FAO pathways are indeed primed and would be more rapidly mobilized in G2 organoid monolayers than their G1 counterparts if provided lipid-enriched media.

Downregulated transcripts in G2 were more difficult to functionally interpret as they group loosely with various facets of innate immune responses. Footprinting and motif analysis suggest restricted binding of AP-1 complex members which have pleiotropic cellular functions that include inflammation and immunity. While intestinal epithelia are not first on the list of major immunity determinants, they do function as the interface with microbial and environmental constituents by coordinating barrier function, luminal sampling by professional immune cells, and secretion of defense molecules. A number of the downregulated genes in G2 are interferon inducible and suggest blunting of type I IFN signaling. A correlation between obesity and repressed type I IFN/antiviral responses has been noted in previous studies using human peripheral blood mononuclear cells(23) and diet-induced obese mice.(24, 25) Thought to arise as a consequence of chronic inflammation in metabolic disease, a dampened ability to mount IFN responses during obesity may also be epigenetically controlled since the transcriptional phenotype persisted in organoid culture and chromatin accessibility was restricted at associated TF binding sites. Interestingly, some degree of overlap in the downregulated G2 and G3 transcripts relative to G1 hint that the innate immune changes are not necessarily linked to the hyperabsorptive phenotype. Future work could evaluate interactions of intestinal organoids in monolayer co-cultures with different immune cell populations to understand the functional implications of the transcriptional profiles associated with each group.

To expand the number of study subjects and evaluate the prognostic potential of specific transcripts as biomarkers, we conducted qRT-PCR in G1 and G2 differentiated organoid monolayers from 20 additional subjects and evaluated the AUC of ΔC_T_ values plotted as receiver-operator curves (ROC). Our sample size (n=10 subjects/group) of G1 and G2 demonstrated statistically significant sensitivity and specificity for 9 of the 10 markers measured. This implies that the G2 phenotype has potential to be characterized through transcriptomic evaluation of intestinal epithelial biopsies in lieu of time- and labor-intensive functional organoid-based assays, and that with further cohort expansion and validation, the development of a qRT-PCR-based diagnostic panel may be feasible.

Organoid cultures also provided evidence that obesity phenotypes are complex and that mechanisms other than intestinal sugar hyperabsorption may contribute to disease. Thus, some obese subject organoids (G3) lacked evidence of altered intestinal absorption and metabolic dysregulation. Profiles of additional donor organoids will be necessary to define the proportion of obese patients with intestinal hyperabsorption, and to understand how this phenotype impacts the severity of their metabolic disease, comorbidities, and potential response to therapeutic intervention. However, this study and prior work(9) highlight the importance of personalized approaches to characterizing metabolic disease phenotypes. It is not surprising that the intestinal epithelium undergoes profound imprinted alterations in the obesogenic state given that prior studies have detected differential DNA methylation patterns in other tissues (blood, liver, pancreas, adipocytes) during obesity in humans(26–28) and in rodents on high-fat diet.(26, 29–32) Future studies to understand drivers of the epigenetic changes and identify strategies to limit or reverse intestinal hyperabsorption in these patients could transform clinical approaches to obesity management.

## METHODS

### Human organoid cultures

Organoid cultures were established from adult jejunal tissue obtained during upper endoscopy procedures or tissue discarded following bariatric surgeries as described previously.(9) Informed consent from study participants was obtained under protocols approved by the Institutional Review Boards at Johns Hopkins University (NA_0038329), Baylor College of Medicine (H-31793 and H-31910), and Mayo Clinic Florida (22-001407). All experiments were performed in accordance with review board guidelines and regulations. Recruited subjects were ≥18 years of age, and tissue samples were de-identified such that only age, sex, and BMI were recorded. Glucose absorption phenotype was determined using differentiated organoid monolayers incubated with 25 mM glucose buffer in the apical compartment for 2 h. Organoids classified as Group 1 were established from non-obese donors (BMI ≤ 29.9 kg/m2) and exhibited low intestinal epithelial glucose absorption (≤125 μM glucose detected in basolateral buffer). Organoids in Group 2 were from obese donors (BMI ≥ 35 kg/m2) and showed elevated intestinal epithelial glucose absorption (>125 μM glucose in basolateral buffer). Organoids in Group 3 were from obese donors (BMI ≥ 35 kg/m2) with glucose absorption profiles that were similar to non-obese donors. BMI classification was based on World Health Organization guidelines. Information regarding patient characteristics (age, sex, BMI, glucose absorption phenotype) is provided in **Table 1**. Organoids were cultured in Expansion Media (EM), as previously described (Advanced Dulbecco’s Modified Eagle Medium/Ham’s F-12, 10 mM HEPES, 0.2 mM GlutaMAX, 50% v/v Wnt3A-conditioned media, 15% v/v R-spondin-1-conditioned media, 10% v/v of Noggin-conditioned media, 50 ng/ml human epidermal growth factor (EGF), 500 nM A83-01, 10 μM SB 202190, 1× B27 supplement, 1 mM N-acetylcysteine).(33, 34) At the time of culture initiation and each weekly culture passage, EM was supplemented with 10 µM CHIR99021 and 10 µM Y-27632 for the first 48 h. Growth factor conditioned media was obtained from cell lines expressing Wnt3A (murine L-Wnt3A, CRL-2647, American Type Culture Collection), R-spondin-1 (human HA-Rspo1-Fc/HEK293, kindly provided by Dr. Calvin Kuo, Stanford University, USA), and Noggin (murine Noggin-Fc/HEK293).(35) EM was refreshed every 2-3 d, and organoids were passaged every 7 days. Passage numbers used in this study ranged from 10 to 81.

### Organoid monolayers

Organoid monolayers were prepared as previously described.(^36, 37^) Briefly, isolated organoid fragments resuspended in EM were plated on 24-well plate permeable tissue culture inserts (Corning Transwell 3470; 0.4 μm pore, 0.33 cm^2^ surface area) coated with collagen IV (10 μg/cm^2^, Sigma C5533). To monitor progress toward confluence, transepithelial electrical resistance (TER) was measured using an EVOM2 voltohmmeter (World Precision Instruments). Monolayers judged to be confluent by TER measurement (>300 Ω·cm^2^) and inspection by light microscopy were transferred to differentiation media (DM; Expansion Media lacking Wnt3A, R-spondin-1, A83-01, and SB 202190). Monolayers were used for experiments 5 days after transfer to DM. For the sequencing experiments, differentiated organoid monolayers were digested with Accutase (MilliporeSigma) at 25 °C for 20 min or until cells exhibited rounding and dissociation by light microscopy. The cell suspension was transferred to a low-binding microcentrifuge tube (USA Scientific) and washed with Sorting Buffer (PBS, 25 mM HEPES pH 7.4, 0.5% bovine serum albumin, 5 mM EDTA). Cells were pelleted at 500 xg, 4 °C, 5 min. Buffer was aspirated and cell pellets were snap frozen in liquid nitrogen for RNA-seq, or resuspended at 75,000 cells/100 µL Anti-Freezing Buffer (Advanced DMEM/F-12, 10% fetal calf serum, 10% DMSO) for ATAC-seq, and immediately stored at −80 °C until processed for the respective sequencing applications.

### Glucose transport

Transport of D-glucose was measured in a procedure similar to that described previously.(9) Differentiated organoid monolayers on Transwell inserts were confirmed to be confluent by TER measurements prior to initiating the experiment. Monolayers were washed once (100 μL upper/apical chamber, 600 μL lower/basolateral chamber) with mannose buffer (50 mM HEPES pH 7.4, 138 mM NaCl, 4.7 mM KCl, 1.25 mM MgSO_4_, 1.25 mM CaCl_2_, 0.5% bovine serum albumin, 5 mM mannose) and then incubated in mannose buffer at 37 °C, 5% CO_2_, 30 min. An aliquot (200 μL) was removed from the lower chamber (time 0 measurement) and replaced with an equivalent volume of fresh mannose buffer. The mannose buffer in the upper chamber was aspirated, replaced with 100 μL of 25 mM glucose buffer (50 mM HEPES pH 7.4, 128 mM NaCl, 4.7 mM KCl, 1.25 mM MgSO_4_, 1.25 mM CaCl_2_, 0.5% bovine serum albumin, 25 mM D-glucose), and then returned to the incubator for 2 h. TER was re-measured after 2 h to confirm that the monolayers remained intact during the incubation, and 200 μL of buffer in the lower chamber was sampled. The glucose concentration in the buffers was measured using an Amplex Red Glucose Assay kit (ThermoFisher Scientific). The final 96-well plate absorbance was read at 570 nm on an AccuSkan microplate reader (ThermoFisher Scientific).

### RNA sequencing

The organoid lines used for RNA-seq and ATAC-seq were previously characterized for glucose absorption phenotype.(9) Total RNA was isolated from cells and purified by DNase I treatment using the RNeasy Micro kit (Qiagen). RNA-seq libraries were prepared by Illumina TruSeq RNA Library Preparation Kit v2 and sequenced on the Illumina HiSeq 4000 platform in the Mayo Clinic Medical Genomics Core. Mayo RNA-sequencing (MAP-RSeq) analysis pipeline was used to perform gene expression analysis. FASTQ readings (paired end; 100 bp) were aligned to hs38 with TopHat2, generating raw gene and exon counts using featureCount, and sample quality was determined by RSeQC. EdgeRv3.8.6 was used for differential expression analysis and to compare groups. Genes with an absolute log_2_ fold change (log_2_FC) >1.5 and false discovery rate (FDR) <0.05 were considered to be significantly differentially expressed.

### ATAC sequencing

Approximately 50,000 nuclei isolated from organoid monolayers were utilized for ATAC-seq. The transposition reaction was carried out in a thermomixer at 37° C for 30 minutes and oscillated at 1,000 rpm. The size of the library DNA was determined by a Fragment Analyzer from the amplified and purified library. The ATAC-seq library was amplified by PCR. By targeting known highly accessible and closed chromatin sites with real-time PCR, the enrichment of each library’s accessible regions was determined and expressed as a fold difference. BWA was used to map paired end reads to the hs38 genome. Picard SortSam was used to convert SAM files into BAM files and sort by chromosomal coordinates. Picard MarkDuplicates was used to remove PCR duplicates. Processed BAM files were used to call the peak by MACS2. BAM files and peak files obtained from MACS2 was processed using DiffBind to identify differentially accessible genomic loci. Using an FDR threshold of 0.05, Homer was used to annotate the differentially accessible loci to the closest transcription start site (TSS). GREAT was used to conduct gene ontology on the genome regions determined in the ATAC-seq analysis with default settings. Motif analysis was completed by the findMotifsGenome.pl command in Homer. Transcription factor footprinting analysis was conducted using the TOBIAS computational framework.(10)

### Quantitative real-time PCR (qRT-PCR)

Organoid lines used for qRT-PCR were newly collected for this study. Total RNA was extracted from differentiated organoid monolayers using the PureLink RNA Mini Kit (Invitrogen) and reverse transcribed to complementary DNA using SuperScript VILO Master Mix (Invitrogen). Reactions for qRT-PCR were carried out in triplicate using 5 ng cDNA, gene-targeted oligonucleotide pairs (sequences in **Table 2**), and Power SYBR Green Master Mix (Invitrogen) on a QuantStudio 12K Flex instrument (Applied Biosystems). For each organoid line, the gene target was normalized to the average C_T_ for 18S ribosomal RNA (RN18S) and GAPDH to obtain ΔC_T_ values. Statistical significance was calculated by unpaired student’s t-test. Receiver-operator curves (ROC) plotting the true positive rate (sensitivity) versus the false positive rate (1 – specificity), and the associated area under the curve and *p*-values for each target, were prepared using GraphPad Prism 10 using 95% confidence intervals and the Wilson/Brown method.

**Table 2.**
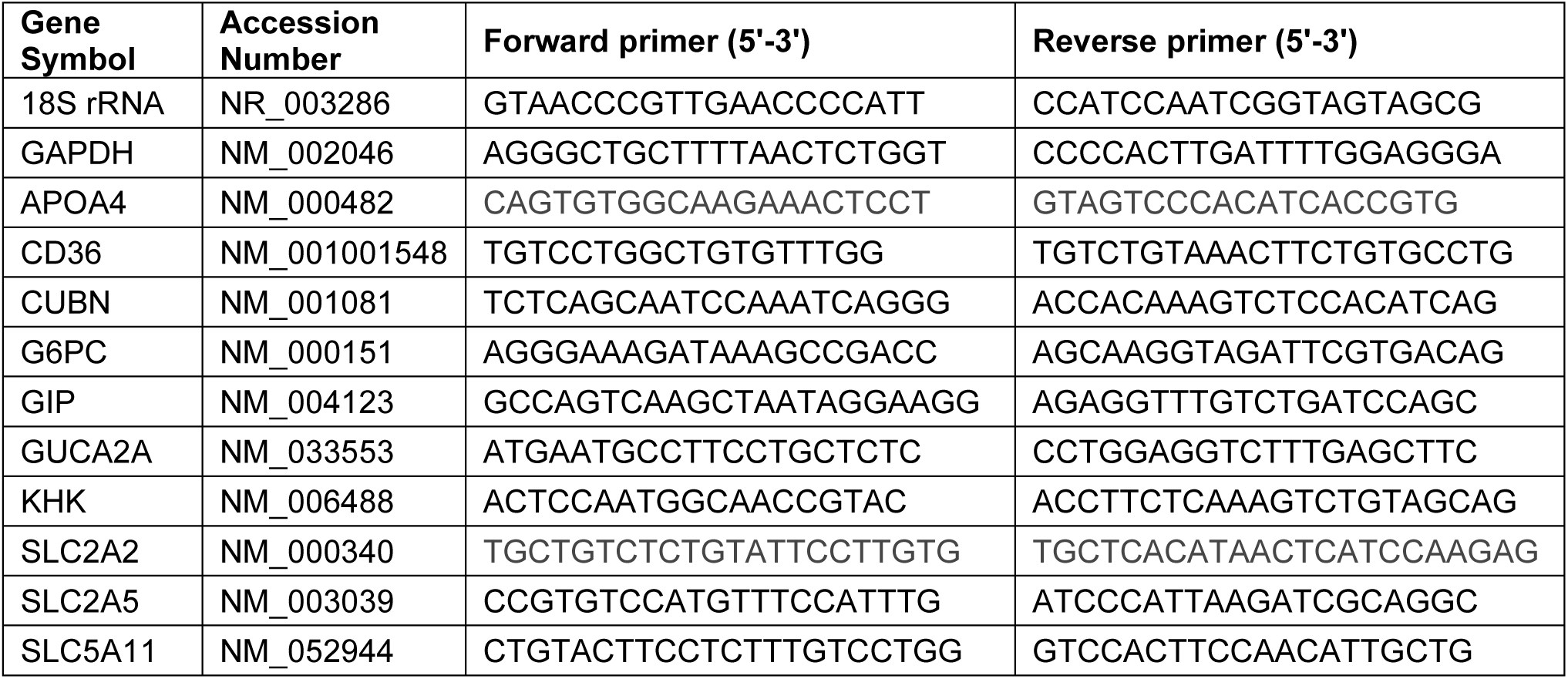
Oligonucleotide sequences. Oligonucleotides used in qRT-PCR assays.

## DATA AVAILABILITY

Data is provided within the manuscript and in supplementary tables. The complete RNA-seq and ATAC-seq datasets generated and analyzed in the current study are available online in the Gene Expression Omnibus (accession number pending).

## ABBREVIATIONS

ATAC-seq: assay for transposase-accessible chromatin by sequencing
AUC: area under the curve
BMI: body mass index
DEG: differentially expressed gene
FDR: false discovery rate
FPKM: fragments per kilobase of transcript per million mapped reads
GOBP: Gene Ontology Biological Process
ISC: intestinal stem cell
qRT-PCR: quantitative real-time polymerase chain reaction
ROC: receiver-operator curves
RYGB: roux-en-Y gastric bypass
TF: transcription factor
TSS: transcriptional start site)

## ACKNOWLEDGEMENTS

The authors thank Dr. Mary K. Estes (Baylor College of Medicine) for sharing four jejunal organoid lines (J2, J4, J11, and J12) used in this study. Tissue culture facilities were provided by the Johns Hopkins Digestive Diseases Basic & Translational Research Core Center (National Institutes of Health P30 DK089502). RNA-seq was performed by the Genome Analysis Core, Mayo Clinic Rochester. ATAC-seq was conducted by the Epigenomics Development Laboratory, Center for Cell Signaling in Gastroenterology, Mayo Clinic Rochester (National Institutes of Health P30 DK084567). Sequence assembly pipelines, differential gene expression, and ATAC footprinting were carried out by the Bioinformatics Core, Mayo Clinic Rochester. Quantitative PCR was conducted on instrumentation provided by the Johns Hopkins Genetic Resources Core Facility.

## DISCLAIMERS

Dr. Kovbasnjuk is an employee of the National Institutes of Health, USA. The content of this manuscript is solely the responsibility of the author and does not necessarily represent the official views of the National Institutes of Health, USA.

## AUTHOR CONTRIBUTIONS

D.S.B., O.K., V.K., and J.F.A. conceived and designed research; D.S.B., Z.L., J.H.L., T.M., A.V.B., R.L., A.D., and J.F.A. performed experiments; D.S.B., Z.L., J.H.L., T.M., A.V.B., T.O., O.K., and J.F.A. interpreted results of experiments; Z.L., T.M., A.V.B, and J.F.A. prepared figures; D.S.B, Z.L., T.O., O.K., and J.F.A. drafted and edited manuscript; all authors approved final version of the manuscript.

